# Sensorimotor conflicts alter perceptual and action monitoring

**DOI:** 10.1101/504274

**Authors:** Nathan Faivre, Laurène Vuillaume, Fosco Bernasconi, Roy Salomon, Olaf Blanke, Axel Cleeremans

## Abstract

Bodily self-consciousness is defined as a set of prereflective representations of integrated bodily signals giving rise to self-identification, self-location and first-person perspective. While bodily self-consciousness is known to modulate perception, little is known about its influence on higher-level cognitive processes. Here, we manipulated bodily self-consciousness by applying sensorimotor conflicts while participants performed a perceptual task followed by confidence judgments. Results showed that sensorimotor conflicts altered perceptual monitoring by decreasing metacognitive performance. In a second experiment, we replicated this finding and extended our results by showing that sensorimotor conflicts also altered action monitoring, as measured implicitly through intentional binding. In a third experiment, we showed that effects on perceptual monitoring were induced specifically by sensorimotor conflicts related to the trunk and not to the hand. Taken together, our results suggest that bodily self-consciousness may serve as a scaffold for perceptual and action monitoring.

## Introduction

The self is a multifaceted construct that minimally entails an organism’s ability to distinguish its constituents from the surrounding environment. It is defined at different levels of complexity (Rochat, 2003), ranging from fundamental biological mechanisms (e.g., homeostasis, immunological tolerance), to bodily representations (e.g., peripersonal space), to more abstract cognitive functions such as self-recognition or autobiographical memory. Here, we sought to describe the interplay between two crucial aspects of the self, and in particular self-consciousness, defined at the bodily and cognitive levels. At the bodily level, self-consciousness includes self-identification (the conscious experience of identifying with the body) and self-location (the experience of where “I” am in space) (for reviews see Blanke & Metzinger, 2009; Blanke, Slater & Serino, 2015; Ehrsson 2012). At the cognitive level, the sense of self includes perceptual monitoring or metacognition, defined as the capacity to monitor and control one’s own perceptual states (Koriat, 2006; Fleming & Frith 2012), and to compute the likelihood of being correct given sensory evidence during perceptual tasks (Pouget, Drugowitsch, & Kepecs, 2016). The cognitive self also includes the capacity to monitor and control one’s own actions, notably to predict the sensory consequences of a motor command (Blakemore and Frith, 2003; Haggard, 2017). The present study aims at assessing the possibility that cognitive functions such as perceptual and action monitoring may rely on multimodal representations relevant to bodily self-consciousness. In support of this view, it was shown that disgust cues modulating bodily reactions like heart rate and pupil dilation also modulate confidence judgments, suggesting that interoceptive bodily signals that are independent of the decisional process can guide metacognition (Allen et al., 2016). Besides interoception, action-related signals were also shown to modulate metacognition: confidence relates to sub-threshold motor activity (Gadjos et al., 2018) and alpha desynchronization over the sensorimotor cortex (Faivre et al., 2018), and is disrupted when transcranial magnetic stimulation pulses are applied to the premotor cortex before or after a visual task disrupt subsequent confidence judgements (Fleming et al., 2015). Plus, metacognitive performance is better for committed vs. observed decisions, suggesting that committing to a decision through a motor action informs confidence (Pereira et al., 2018). Together, these studies suggest that interoceptive and action-related signals from the body may play a role for metacognition.

Here, we sought to investigate the interplay between bodily self-consciousness and cognitive functions by measuring the quality of perceptual monitoring in healthy subjects while their bodily representation was systematically manipulated through the application of sensorimotor conflicts. Participants were asked to perform tapping movements with a robotic device situated in front of them, while another robot connected to the front device applied corresponding tactile stimuli on their back (synchronous condition). In the asynchronous condition, a constant temporal delay between the movement of the participant and the tactile stimulation delivered by the back robot was introduced, which has the effect of increasing prediction errors regarding the sensory consequences of a motor command. Such manipulations are also known to induce alterations of bodily self-consciousness such as changes in self-location (Blanke et al., 2014). Assuming that the mechanisms enabling perceptual and action monitoring relate to those enabling bodily self-consciousness, we expected alterations of self-location induced by sensorimotor conflicts to induce impairments of perceptual and action monitoring. In Experiment 1, we quantified the capacity of participants to monitor their performance on an auditory temporal order judgment task while actuating the robot synchronously or asynchronously. Experiment 2 aimed at replicating the results found in Experiment 1 with a new group of participants, and further quantified their capacity to monitor action consequences during the synchronous vs. asynchronous condition. Finally, Experiment 3 aimed at determining whether effects on perceptual and action monitoring were specific to sensorimotor conflicts impacting full-body representations (Blanke et al., 2014), or whether they could also be induced by similar conflicts impacting limb-representations only. Together, these three experiments show that perceptual monitoring is altered by sensorimotor conflicts centered on the trunk impacting full-body representations, while action monitoring is altered by sensorimotor conflicts impacting both full-body and limb representations. This indicates that bodily-self consciousness may serve as a scaffold for complex cognitive functions including perceptual and action monitoring.

## Method

### Participants

A total of 54 different participants were recruited: 18 in Experiment 1 (10 females, mean age 22.7 years, SD 4.5 years), 18 in Experiment 2 (12 females, mean age 23.7 years, SD 4.2 years) and 18 in Experiment 3 (12 females, mean age 24.1 years, SD 4.2 years). Two participants had to be excluded due to a technical issue during data recording (one in Experiment 1 and one in Experiment 2) as they could not perform the temporal order judgment task). All participants were right-handed, had normal hearing and no psychiatric or neurological history, and participated in exchange for a monetary compensation (20 CHF per hour). They were naive to the purpose of the study and gave informed consent, in accordance with institutional guidelines and the Declaration of Helsinki. The study was approved by the cantonal ethics committee in Geneva. The sample size in Experiment 1 was predefined based on a pilot study, and was kept constant in Experiment 2 and 3.

### Apparatus and stimuli

Robotic System: we used a system composed of a commercial haptic interface (Phantom Omni, SensAble Technologies), coupled with a three degree-of-freedom robot in the back (see Fig. 1 and Hara et al., 2011; Blanke et al., 2014 for details). Participants were standing and controlling the front robot situated directly in front of them with their right index finger (excepted in the baseline condition of Experiment 1 in which it was controlled by the experimenter). The back robot was placed directly behind their back and reproduced with virtually no delay the movements produced with the front robot in the synchronous condition, and with 500 ms delay in the asynchronous condition. Participants were asked to perform tapping movements in every direction to touch their back on a 200 mm × 250 mm surface. In Experiment 3, the same setup was used except that the back robot was adjusted to point in the vertical axis so to touch the participants hand instead of their back. Participants could again perform any tapping movements they wanted as long as the robot touched the back of their hand.

**Figure 1:**
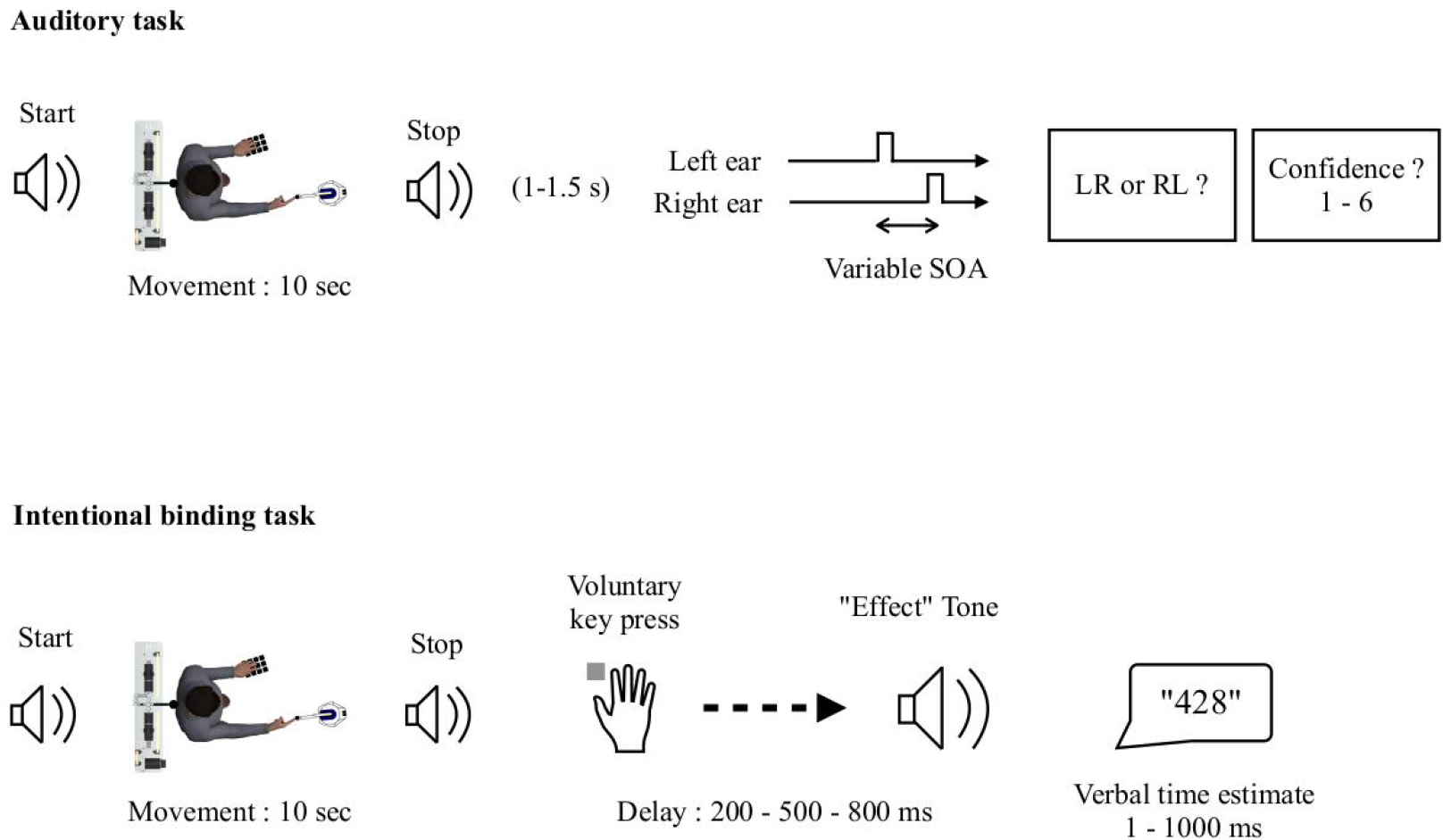
Experimental procedure. After actuating the front robot and receiving synchronous or asynchronous tactile feedback for 10 s, participants were asked to perform one of two tasks. In the auditory task (upper row) participants had to indicate whether they heard a sequence of two sounds starting in the left and ending in the right ear or vice versa (i.e., temporal order judgment task). They were then asked to report how confident they were in their response. Both responses were given using the left hand. In the intentional binding task (lower row) participants were asked to press a key with the left hand, and report verbally the delay with which a subsequent effect tone was played.

Auditory stimuli: all experimental sounds were sinusoidal pure tones, with 1 ms rise/fall time and 44100 Hz sampling rate, generated using MATLAB (MathWorks, Natick, MA) with the Psychophysics toolbox (Brainard, 1997; Pelli, 1997; Kleiner, Brainard, Pelli 2007). Auditory stimuli used for the temporal order judgement task were 600 Hz pitch pairs of sounds, played for 10 ms via headphones either to the left and then to the right ear (Left–Right or LR) or to the right and then to the left ear (Right–Left or RL), with a variable stimulus onset asynchrony (SOA) that was adjusted throughout the experiment using an adaptive one-up two-down staircase procedure (Levitt, 1971). The initial SOA was set to 80 ms, and varied in 5 ms steps between 5 ms and 150 ms. Cue sounds (400 Hz pitch, 100 ms duration,) served as indicators of the beginning and the end of each trial. White noise was played in both ears during the whole experiment to isolate the participant from external noises. The sound pressure level was adjusted before the experiment individually at a comfortable level with the auditory stimuli volume always four times higher than the white noise volume.

### Procedure

Prior to the experiment, participants were told about the general experimental procedure, and were instructed in the use of the robot. After filling in a questionnaire for demographic data, participants were equipped with headphones and blindfolded. While standing, they were asked to insert their right index finger into the front device and perform tapping movements, which lead the back robot to deliver tactile pokes on their back. They were allowed to move the front device in any direction along the vertical and horizontal axes, which resulted in pokes applied to different parts of their back. The main task was as follows: each trial started with a cue sound indicating to start the tapping movements with the right index finger. After 10 s of tapping, a second cue sound was played, indicating to stop moving. Following a random interval between 1000 and 1500 ms duration, participants were presented with two successive sounds and were asked to indicate by means of keypress with the left hand whether they perceived an LR or RL pair (temporal order judgment, Bernasconi et al., 2010). This first response defined performance for the first order task, for which no feedback was provided. Subsequently, as a second-order task, participants were asked to report the confidence they had in their response by pressing a key with their left hand between 1 (very unsure) to 6 (very sure). A random inter-trial interval between 1000 and 1500 ms was enforced. The experiment contained three main conditions grouped in blocks. In the synchronous condition, the back device responded to the front robot actuated by the participants with virtually no temporal delay (Hara et al., 2011). In the asynchronous condition, a delay of 500 ms was set between the front and the back devices, so that participants felt a poke on their back 500 ms after moving the front device. The asynchronous condition resulted in a spatiotemporal sensorimotor conflict between the right hand actuating the front robot and the back receiving tactile feedback. Such condition is known to induce global changes in bodily self-consciousness, notably in terms of self-location (Blanke et al., 2014). In the baseline condition, participants passively received tactile feedback while the front robot was actuated by the experimenter. While actuating the front robot in the synchronous and asynchronous conditions, participants received a somatosensory force feedback on their right index finger each time the back robot touched their back, so to mimic the effect of physical resistance. The experiment was divided in blocks of 30 consecutive trials of the same condition, with a total of 9 blocks (3 in succession per condition) counterbalanced across participants. A training phase of 12 trials was enforced before starting the experiment. At the end of the first block of each condition, participants were asked to fill in a questionnaire composed of 10 Likert scale items: 1) I felt as if I had no body. 2) I felt as if I was touching my body. 3) I felt as if I was touching someone else’s body. 4) I felt as if I was in front of my body. 5) I felt as if I was behind my body. 6) I felt as if I had more than one body. 7) I felt as if someone else was touching my body. 8) I felt as if I was touched by a robot. 9) I felt as if someone was standing behind my body. 10) I felt as if someone was standing in front my body. The experiment lasted 120 minutes and ended with an individual debriefing.

Experiment 2 was divided into two sessions. The first session followed the exact same procedure as Experiment 1 (i.e., first and second-order tasks), except that it contained no baseline condition, and therefore lasted 80 min instead of 120 min. The second session relied on the classical intentional binding task (Haggard, Clark & Kalogeras, 2002; Wenke & Haggard, 2009), in which participants were asked to press a key with their left hand whenever they felt the urge to do so. The keypress triggered a target tone (600 Hz pitch, 200 ms duration) after a temporal delay of 200 ms, 500 ms or 800 ms. Participants were told that the target tone could occur after a random delay between 1 ms and 1000 ms following key press, and were asked to report verbally their best estimate for this delay. After reporting their estimate, they had to press a key to start the next trial. Participants were actuating the front robot with their right hand for the entire trial duration. Session 2 contained a synchronous and asynchronous condition like session 1. Participants completed two blocks of 30 trials per condition, corresponding to 10 repetitions for each temporal delay. The order of conditions was counterbalanced across participants, and remained identical within participant for sessions 1 and 2. The order of temporal delays was randomized across trials. A training phase of 12 trials was enforced before starting session 2. It ended with an individual debriefing and its total duration was about 70 min. A break of 30 min was allowed between session 1 and 2. At the end of session 2, participants were asked to actuate the robot for 1 minute (Synchronous and Asynchronous in the same order as in session 1 and 2), and then filled in the same questionnaires as in Experiment 1 (see below). This was performed at the very end of the experiment to avoid demand characteristics effects (Orne, 1962).

Experiment 3 was identical to Experiment 2, except that participants were seated and that the stroking was applied on the back of their left hand instead of on their back.

### Questionnaire

Participants were asked to rate specific aspects of the subjective experience they had in the different experimental conditions. The questions were based on a previous study (Blanke et al., 2014, see supplementary data) and investigated in particular the subjective feeling of touching oneself (“I felt as if I was touching my body”; self-touch) or of touching somebody else’s body (“I felt as if I was touching someone else’s body”; other-touch). Other questions investigated the subjective sensation of corporeal displacement (i.e. “I felt as if I was in front of my body”) and the feeling of a presence (i.e. “I felt as if someone was standing behind my body.”). Other items served as control questions for suggestibility (i.e. “I felt as if I had no body”). Ratings were reported on a Likert scale from 0 (Not at all) to 6 (Very strong) and transformed into Z-scores prior to statistical analysis.

### Data analysis

Reaction times for temporal order judgments longer than 3 s and shorter than 300 ms were discarded (corresponding to 6.2 % of total trials in Experiment 1, 6.4 % in Experiment 2, and 11.4 % in Experiment 3). Reaction times for confidence judgements longer than 6 s and shorter than 300 ms were discarded (corresponding to 3.0 % of total trials in Experiment 1, 2.0 % in Experiment 2, and 4.7 % in Experiment 3).

Metacognitive performance was analysed with two different approaches. First, we performed mixed effects logistic regressions between accuracy and confidence, and considered the regression slope as an indicator of metacognitive performance (that is, the capacity for a participant to adapt confidence to performance), and the lower asymptote as a measure of confidence bias (that is, the capacity to report low confidence estimates when perceptual evidence is low). This approach is agnostic regarding the signals used to compute confidence estimates (i.e., decisional vs. post-decisional locus, see Yeung & Summerfield, 2012; Pleskac & Busemeyer, 2011), and the mixed model framework allows analysing raw confidence ratings even if they are unbalanced (e.g., in case participants do not use all possible ratings) (Rausch et al., 2015). Second, relying on signal detection theory, we quantified metacognitive sensitivity with meta-d’ (Maniscalco & Lau, 2012, 2014), which reflects the amount of perceptual evidence available when performing confidence judgments. Contrary to the logistic regression approach, signal detection theory assumes that confidence judgments are informed by perceptual evidence only, with no contribution of post-decisional processes. The resulting measure of metacognitive sensitivity (meta-d’) shares the same dimension as perceptual sensitivity (d’), which allows normalizing one by the other, and deriving an index of metacognitive performance independent of task performance, called metacognitive efficiency (meta-d’/d’). Meta-d’ was computed following a resampling of confidence ratings: for a given participant and condition, confidence ratings used in less than 10 trials were merged with the superior rating (e.g., if one participant gave a confidence rating of 1 in 6 trials, and of 2 in 18 trials, we merged the two categories in 24 trials with a confidence rating of 2). This ensured that the fit by maximum likelihood estimation involved in the computation of meta-d’ was performed on a sufficient number of points (Maniscalco & Lau, 2012, implemented in R by Rausch et al., 2015). The tendency to report high or low confidence ratings independently of task performance was quantified with confidence bias, based on the type 2 receiver operating characteristic curve (ROC) which determines the rate of correct and incorrect responses at each confidence level. Specifically, the area between the ROC and major diagonal was divided by the minor diagonal, and confidence bias was defined as the log ratio of the lower and upper area (Kornbrot, 2006).

Response times in the intentional binding task were analysed using linear mixed effects regressions, with condition and delay as fixed effects, intercepts for subjects as random effects, and a by-subject random slope for the effect of condition and delay. Reaction times below or above 2 standard deviations away from the mean were discarded for each subject and each delay (corresponding respectively to 3.7% and 4.2% of total trials in Experiment 2 and 3). As response times were not normally distributed, they were considered as ordinal data and rank-transformed before linear mixed modelling (Conover & Iman, 1981). All analyses were performed with R (2016), using notably the afex (Singmann et al., 2015), BayesFactor (Morey et al., 2015), ggplot2 (Wikham, 2009), lme4 (Bates et al., 2014), lmerTest (Kuznetsova, Brockhoff & Christensen, 2015), and effects (Fox, 2003) packages. In all ANOVAs, degrees of freedom were corrected using the Greenhouse-Geisser method.

## Results

### Perceptual monitoring

#### Experiment 1

Regarding the first-order task (temporal order judgment), an analysis of variance revealed that the SOA corresponding to perceptual threshold differed across conditions (F(1.83,27.39) = 8.02, p = 0.002, η_p_^2^ = 0.35), with lower SOA in the baseline (mean SOA = 45 ms, SD = 13 ms) vs. synchronous condition (mean SOA = 53 ms, SD = 14 ms; paired t-test: p = 0.020) and in the baseline vs. asynchronous condition (mean SOA = 56 ms, SD = 15 ms; paired t-test: p < 0.001), but no difference between the synchronous and asynchronous conditions (paired t-test: p = 0.36, BF = 0.37). This implies that the task was easier in the baseline compared to the synchronous and asynchronous conditions, which is expected considering that participants performed no tapping movement in the baseline condition. Despite these differences in terms of task difficulty, task performance was equated with the staircase procedure we used (Levitt, 1971), and no effect of condition on sensitivity (d’: F(1.65,24.78) = 0.93, p = 0.39, η_p_^2^ = 0.06), criterion (F(1.56,23.39) = 0.74, p = 0.46, η_p_^2^ = 0.05), or reaction times (F(1.78,26.71) = 1.48 p = 0.24, η_p_^2^ = 0.09) was found, revealing that task performance was adequately controlled across conditions. Regarding the second order task, we found no effect of condition on raw confidence ratings (F(1.94,29.16) = 1.12, p = 0.34, η_p_^2^ = 0.07), confidence bias (F(1.65,24.68) = 2.4, p = 0.12, η_p_^2^ = 0.14), or reaction times for providing confidence ratings (F(1.96,29.37) = 0.65, p = 0.53, η_p_^2^ = 0.04), revealing that the production of confidence estimates per se was not impacted by our manipulation.

Next, we assessed how confidence ratings tracked first order accuracy, by fitting a mixed effects logistic regression on task accuracy, with condition and confidence as fixed effects, intercept for participants as random effects, and a by-subject random slope for the effect of confidence. First, the model revealed higher intercepts in the asynchronous compared to the baseline condition (estimate = 0.46, Z = 1.99, p = 0.047), and similar intercepts between the baseline and the synchronous condition (estimate = −0.12, Z = −0.12, p = 0.60). This indicates that in the asynchronous condition participants had a higher first-order accuracy when reporting guessing than in the synchronous and baseline conditions. Crucially, the model revealed that the relation between confidence and accuracy differed in the asynchronous vs. baseline condition (estimate = −0.16, Z = −2.48, p = 0.013), but not between the synchronous and baseline condition (estimate = −0.02, Z = −0.26, p = 0.80). As can be seen on Fig. 2, this is reflected by a slope of smaller magnitude in the asynchronous compared to the synchronous and baseline conditions, which indicates a decrease in the capacity to adapt confidence to task performance, while task performance was similar across conditions. Importantly, this effect on metacognitive performance cannot be explained by the difference in SOA reported above, as no slope difference was found between the synchronous and baseline conditions, while SOA differed between these two conditions. Plus, another mixed effects logistic regression comparing only the synchronous and asynchronous conditions revealed different intercepts (estimate = 0.55, Z = 2.36, p = 0.018) and slopes (estimate = −0.14, Z = −2.16, p = 0.031), confirming that metacognitive performance was lower in the asynchronous vs. synchronous conditions, this despite an equal SOA between the two conditions. We conclude that a specific decrease in metacognitive performance occurred in the asynchronous condition.

**Figure 2:**
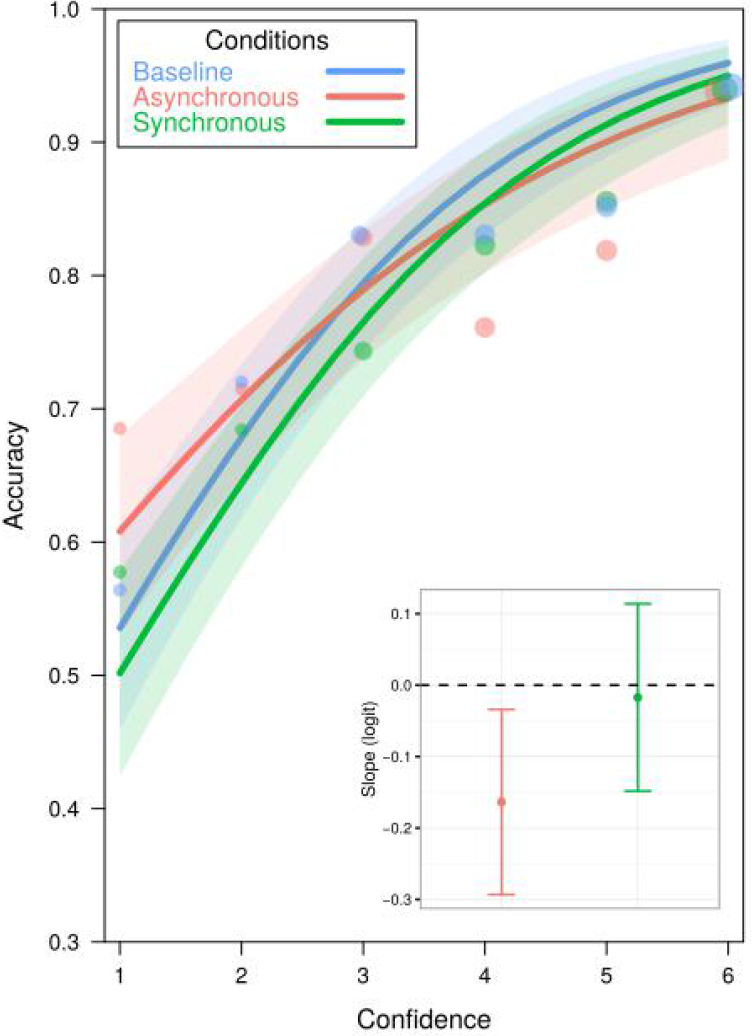
Mixed logistic regression between task accuracy and confidence in the baseline (blue), asynchronous (red), and synchronous condition (green) in Experiment 1. Each dot represents the group-average accuracy for a given level of confidence, with dot size representing the number of total trials in that specific condition. The shaded area around each fit represents the 95% confidence interval. The inset plot represents the estimated slope in logit unit in the asynchronous (red) and synchronous (green) conditions, with respect to the baseline condition (horizontal dashed line). Error bars represent the 95% confidence interval.

#### Experiment 2

We then sought to replicate these findings in Experiment 2. Compared to Experiment 1, a direct comparison between the synchronous and asynchronous conditions was performed, with no additional baseline. Analyses of variance revealed no difference in task performance for the temporal order judgments between the synchronous condition and the asynchronous condition. There was no effect of condition on SOA (F(1,16) = 1.35, p = 0.26, η_p_^2^ = 0.08), sensitivity (F(1,16) = 0.02, p = 0.88, η_p_^2^ = 0.00), criterion (F(1,16) = 0.88, p = 0.36, η_p_^2^ = 0.05), or reaction times (F(1,16) = 2.96, p = 0.10, η_p_^2^ = 0.16).

Regarding confidence ratings, we found no effect of condition on confidence (F(1,16) = 0.47, p = 0.50, η_p_^2^ = 0.03), confidence bias (F(1,16) = 0.37, p = 0.55, η_p_^2^ = 0.02), or reaction times for confidence ratings (F(1,16) = 3.12, p = 0.10, η_p_^2^ = 0.16). The same mixed effects logistic regression as in Experiment 1 was then used to assess how confidence ratings tracked first order accuracy. The model revealed similar intercepts between the synchronous and the asynchronous conditions (z = −1.57, p = 0.12) and an effect of condition on the relation between confidence and accuracy (z = −2.05, p = 0.040) (see Fig. 3). Similarly to Experiment 1, this indicates a decrease in metacognitive performance in the asynchronous condition independently of any change in task performance.

**Figure 3:**
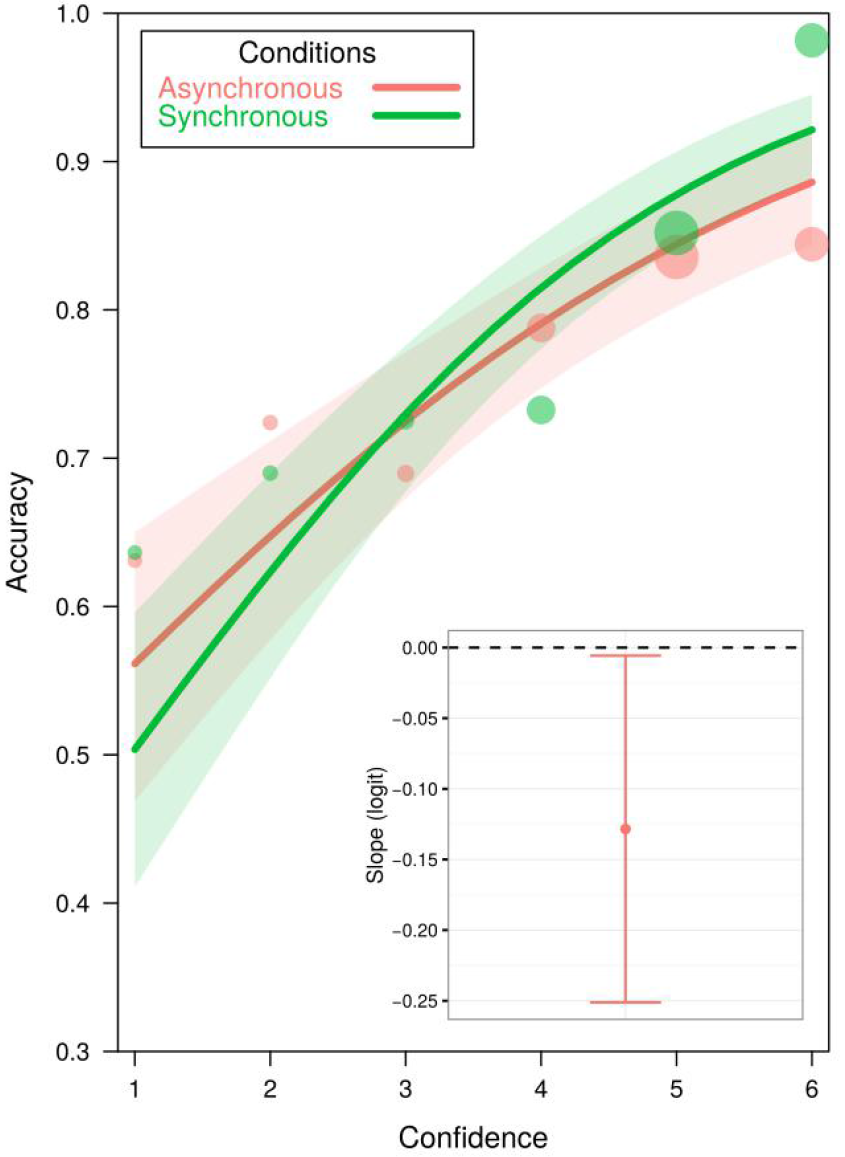
Mixed effects logistic regression between task accuracy and confidence in the asynchronous (red), and synchronous condition (green) in Experiment 2. Each dot represents the group-average accuracy for a given level of confidence, with dot size representing the number of total trials in that specific condition. The shaded area around each fit represents the 95% confidence interval. The inset plot represents the estimated slope in logit unit in the asynchronous vs. synchronous condition (horizontal dashed line). Error bars represent the 95% confidence interval.

As an alternative to logistic regressions, we attempted to replicate our findings relying on signal detection theory to assess metacognitive performance. Specifically, we used the ratio of meta-d’/d’ as an index of metacognitive efficiency, that is the amount of perceptual evidence available to perform confidence judgements. Lower metacognitive efficiency in the asynchronous vs. synchronous condition was confirmed in Experiment 1 (one-tailed paired t-test: t(15) = 2.21, p = 0.02) and in Experiment 2 (one-tailed paired t-test: t(16) = 1.88, p = 0.04) (Figure 4). These results based on signal detection theory confirm our previous results that metacognition is altered in the presence of sensorimotor conflicts, and rule out any possible confound in terms of first-order task performance.

**Figure 4:**
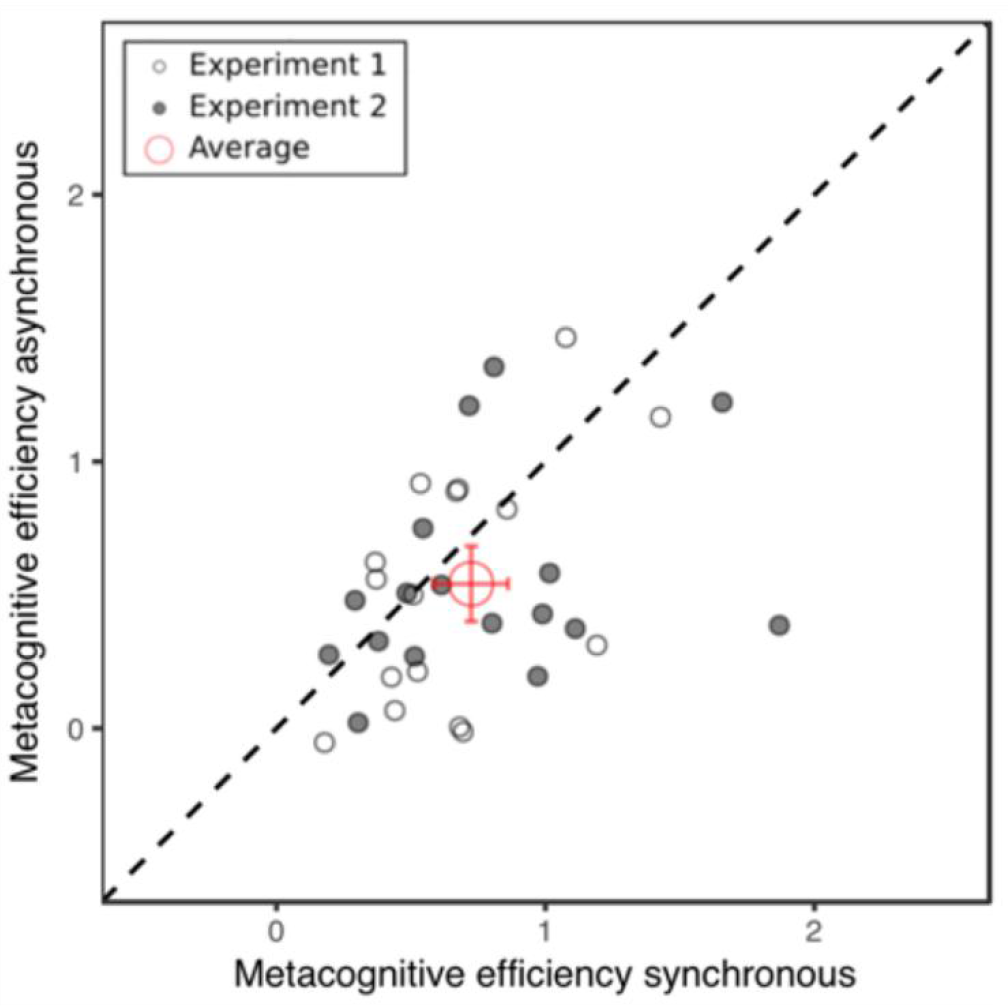
Metacognitive efficiency in the asynchronous vs. synchronous condition for each participant in Experiment 1 (empty dots) and 2 (full dots). Dots lying below the diagonal reflect lower metacognitive sensitivity in the asynchronous condition. The red dot corresponds to the average across all participants, error bars represent 95% confidence interval.

#### Experiment 3

To further define the nature of sensorimotor conflicts susceptible of altering metacognition, we ran a third experiment identical to Experiment 2, except that the back robot touched the left hand instead of the trunk, thereby inducing a more local, hand-related, sensorimotor conflict between the right hand actuating the front robot and the left hand receiving tactile feedback (see Blanke 2012, 2015 for the link between hand/trunk representations and bodily self-consciousness). Following the same analysis strategy, we first ran an ANOVA on participant’s temporal order judgments which revealed no difference in task performance. There was no effect of condition on SOA (F(1,17) = 4.02, p = 0.06, η_p_^2^ = 0.19), first order sensitivity (F(1,17) = 0.27, p = 0.61, η_p_^2^ = 0.02), criterion (F(1,17) = 0.27, p = 0.61, η_p_^2^ = 0.02) or reaction times (F(1,17) = 0.95, p = 0.34, η_p_^2^ = 0.05).

There was no effect of condition on raw confidence ratings (F(1,17) = 0.3, p = 0.59, η_p_^2^ = 0.02), confidence bias (F(1,17) = 1.29, p = 0.27, η_p_^2^ = 0.07), or reaction times for confidence ratings (F(1,17) = 0.3, p = 0.59, η_p_^2^ = 0.02). To assess how confidence ratings tracked first order accuracy, the same mixed effects logistic regression as in Experiment 1 and 2 was used. It revealed similar intercepts (z = −0.94, p = 0.35) and similar slopes (z = 1.19, p = 0.23) between the synchronous and the asynchronous conditions (see Fig. 5). Likewise, metacognitive efficiency did not differ across conditions (F(1,17) = 0.2, p = 0.66, η_p_^2^ = 0.01, BF = 0.27). This indicates that metacognitive monitoring was not impacted when similar sensorimotor conflicts altered limb-based representation instead of trunk-based body representation.

**Figure 5:**
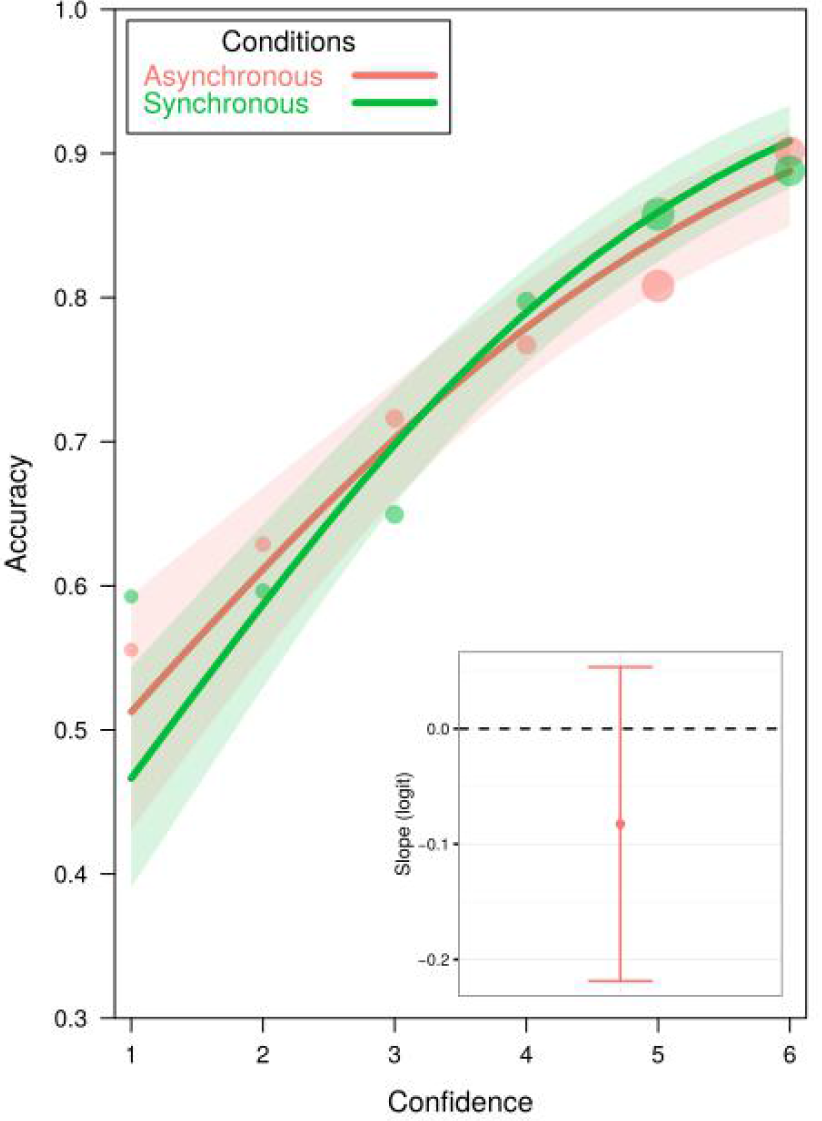
Mixed effects logistic regression between task accuracy and confidence in the asynchronous (red), and synchronous condition (green) in Experiment 3. Each dot represents the group-average accuracy for a given level of confidence, with dot size representing the number of total trials in that specific condition. The shaded area around each fit represents the 95% confidence interval. The inset plot represents the estimated slope in logit unit in the asynchronous vs. synchronous condition (horizontal dashed line). Error bars represent the 95% confidence interval.

### Action monitoring

In addition to perceptual monitoring, we examined the link between bodily self-consciousness and conscious action monitoring, commonly referred to as the sense of agency (Blakemore and Frith, 2003; Gallagher, 2000; Moore and Obhi, 2012). The sense of agency was quantified using intentional binding (Haggard, Clark, Kalogeras, 2002), an implicit measure in which participants have been shown to underestimate the delay between a voluntary action and its consequence. Here, while actuating the front device with the right hand, participants were asked to press a button with their left hand whenever they felt the urge to do so, and had to estimate the delay between this key press and the onset of a sound played 200, 500, or 800 ms after. In experiment 2, a linear mixed effects on ranked response times revealed no main effect of condition (F(1,16.01) = 2.85, p = 0.11), but a main effect of delay (F(2,15.99) = 93.57, p < 0.001), showing that participants reported longer durations when the delay between their key press and the sound onset increased. More importantly, the model revealed a significant interaction between delay and condition (F(2,1888.48) = 3.96, p < 0.02), indicating that participants judged the intervals as significantly shorter in the asynchronous vs synchronous condition, and that this effect was present mainly for long delay (see Fig. 6, left panel). In other words, we found a relative compression of time between a voluntary action and its outcome, if participants were receiving additional asynchronous vs synchronous sensorimotor stimulation.

**Figure 6:**
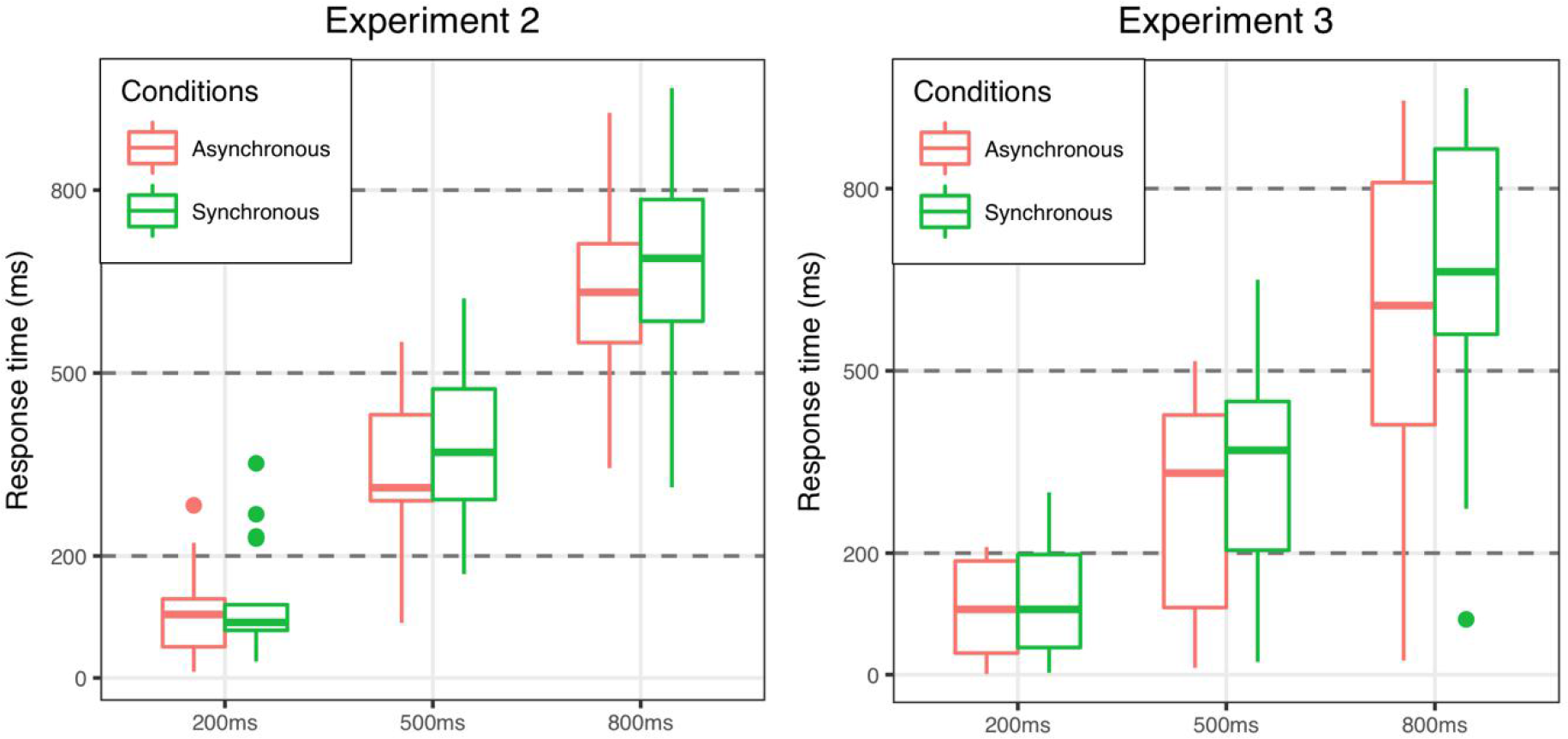
boxplots of estimated response times as a function of delay in the asynchronous (in red) and synchronous (in green) conditions in Experiment 2 (left panel) and Experiment 3 (right panel).

The same analysis confirmed these results in Experiment 3, where participants actuated the front robot with their right hand, received tactile feedback on their left hand, and used the left hand to press a key whenever they felt the urge to do so. We found a main effect of delay (F(2,17.28) = 90.23, p < 0.001), indicating again that participants adapted their response as a function of the delay, and a main effect of condition (F(1,15.05) = 11.81, p < 0.004), showing that participants reported overall shorter times in the asynchronous vs. synchronous conditions (i.e., intentional binding). As in Experiment 2, a significant interaction between condition and delay (F(2,1782.10) = 5.76, p < 0.004) indicated that this effect was more pronounced at longer delays (see Fig. 6, right panel).

### Questionnaire results

Regarding the questionnaire results in the 3 experiments we found that participants felt as if they were touching their own body as significantly higher in the synchronous condition (mean = 2.58, SD = 1.94 for Experiments 1 and 2 and mean = 4.44, SD = 1.15 for Experiment 3) than in the asynchronous condition (mean = 1.48, SD = 1.30 for Experiments 1 and 2 and mean = 2.72, SD = 1.71 for Experiment 3; F(1,32) = 13.36, p < 0.001, η_p_^2^ = 0.29 for Experiments 1 and 2 combined and F(1,17) = 24.53, p < 0.001, η_p_^2^ = 0.59 for Experiment 3). Participants also reported a forward-drift in self-location in the synchronous condition (mean = 1.12, SD = 1.56) compared to the asynchronous condition (mean = 0.97, SD = 1.61) for Experiments 1 and 2 (F(1,32) = 7.49, p = 0.01, η_p_^2^ = 0.19). No other questions were found significantly different between conditions.

## Discussion

With three independent experiments, we examined the influence of sensorimotor conflicts known to alter bodily self-consciousness on two distinct cognitive functions, namely perceptual and action monitoring. While sensorimotor conflicts were induced between the right hand and back (Experiments 1 and 2) or between the right hand and left hand (Experiment 3), we asked participants to estimate the confidence they had regarding their performance on a concurrent auditory task (i.e., perceptual monitoring), or to estimate the delay between a keypress they made spontaneously and an auditory cue (i.e., action monitoring). These two measures served as a proxy to quantify metacognitive performance and intentional binding, respectively.

### Bodily self-consciousness and perceptual monitoring

Regarding metacognitive performance, mixed effects logistic regression analyses showed that when receiving asynchronous sensorimotor feedback on their back, participants were less able to adjust their confidence to performance, and overperformed when reporting guessing. This indicates that sensorimotor conflicts which are known to modulate bodily self-consciousness also impair perceptual monitoring. We replicated these results in a new independent group of participants, and ruled out several experimental confounds. First, the possibility that this decrease in metacognitive performance derived from differences at the perceptual level was excluded by equating first-order performance across conditions, and by re-analysing confidence judgments with a signal detection theory approach which accounts for potential differences in first-order performance (Maniscalco & Lau, 2012). Of note, this approach assumes that confidence estimates are computed based on the same evidence as the perceptual task, while the mixed effects logistic regression approach assumes that confidence can be based both on decisional and post-decisional cues (see Pereira et al., 2018 for recent results disentangling decisional and post-decisional contributions to confidence). As metacognitive impairments were found relying on signal detection theory and mixed logistic regression approaches, we cannot determine whether they have a decisional or post-decisional origin. Second, it is unlikely that participants performed poorly in the asynchronous condition simply due to tactile stimuli they could not predict based on their motor behaviour (i.e., attentional capture). Indeed, we measured similar metacognitive performance in the baseline condition, in which participants passively received tactile stimulation without having to move their right arm to actuate the front robot. Therefore, we argue that this decrease in metacognitive monitoring is neither inherent to deficits at the perceptual level nor due to attentional capture, but rather that it stems from the full-body sensorimotor conflict. It should be noted however that there was no correlation between the questionnaire ratings and the metacognitive deficit, suggesting that it was indeed the sensorimotor mismatch in the asynchronous condition that was important and not necessarily a modulation of bodily self-consciousness as assessed by verbal questionnaires. Interestingly, this specific decrease in metacognitive monitoring did not occur when the same sensorimotor conflicts were applied on the participants’ hands rather than the back. This null result was corroborated by Bayesian analyses supporting the null hypothesis. The role of sensorimotor processing for perceptual monitoring has been a topic of resent research, notably with studies showing a role of motor actions for confidence (e.g., Siedlecka, Paulewicz, & Wierzchoń 2016; Gadjos et al., 2018; Faivre et al., 2018; Pereira et al., 2018). The present study is the first pointing at the specificity of trunk-related signals and bodily-self consciousness for perceptual monitoring.

### Bodily-self consciousness and action monitoring

We also estimated how the same sensorimotor conflicts mentioned above modulated another component of the sense of self, namely the capacity to monitor one’s actions. As a implicit measure, we used intentional binding, defined as the underestimation of the delay between a voluntary action and its consequence (Haggard et al., 2002; Wenke & Haggard, 2009). In two experiments, we measured that intentional binding was stronger in the asynchronous vs synchronous condition, indicating that when participants were exposed to asynchronous sensorimotor conflicts, they perceived actions that were not immediately followed by consequences as their own. This suggests that they monitored the consequences of their actions less accurately in the presence of sensorimotor conflicts known to alter the way they represent their body. As opposed to what we observed for perceptual monitoring, intentional binding was increased both when sensorimotor conflicts were applied to the trunk or to the hand, suggesting that this effect was not specific to full-body manipulations, but rather to the sensorimotor conflict per se, reminiscent of dynamic temporal recalibrations in sensorimotor pathways (Stetson et al., 2006). The directionality of this effect (i.e., more binding in asynchronous vs. synchronous condition) remains to be further explored. One potential issue here is that the dependent variable (i.e., (a)synchrony between an action performed with the left hand and its auditory consequence) was closely related to the manipulation (i.e., (a)synchrony between an action performed with the right hand and its tactile consequence). Therefore, one possibility is that the observed differences of intentional binding may reflect differences in temporal processing unspecific to action monitoring. Future experiments altering the bodily self with other means than asynchronous multisensory conflicts will allow disentangling these two aspects.

## Conclusion

Together, our results extend the recent studies documenting the impact of the bodily self on low-level vision (Faivre et al., 2017; see Faivre, Salomon & Blanke, 2015 for review), and semantic processing of words (Canzoneri et al., 2016; Noel, Blanke, Serino & Salomon, 2017), by further showing that the bodily self may serve as a scaffold for high-level mental capacities which enable the monitoring of one’s thoughts and actions. This is broadly consistent with the idea that there exist deep interactive loops between the self, metacognition and perceptual awareness (Cleeremans, 2011; Timmermans, Schilbach, Pasquali & Cleeremans, 2012), an hypothesis that is at the core of Cleeremans’ Radical Plasticity Thesis.

## Acknowledgments

This work was supported by Swiss National Science Foundation 51AU40_125759, the Bertarelli Foundation and a European Research Council Advanced Grant RADICAL to A.C. N.F. was an Ecole Polytechnique Fédérale de Lausanne Fellow cofunded by Marie Skłodowska-Curie. R.S. was supported by the National Center of Competence in Research (SYNAPSY: The Synaptic Bases of Mental Diseases), financed by Swiss National Science Foundation 51AU40_125759. A.C. is a Research Director with the F.R.S.-FNRS (Belgium).

**Author contribution:** NF, AC, and OB developed the study concept. NF, LV, FB contributed to the study design. Testing and data collection were performed by LV. NF and LV performed data analysis. NF and LV drafted the paper, and all authors provided critical revisions. AC is a research director with the F.R.S.-FNRS. This work was supported by ERC Advanced Grant RADICAL to AC. All authors approved the final version of the paper for submission.

## References

Allen, M., Frank, D., Schwarzkopf, D. S., Fardo, F., Winston, J. S., Hauser, T. U., & Rees, G. (2016). Unexpected arousal modulates the influence of sensory noise on confidence. Elife, 5, e18103.

Bates, D., Mächler, M., Bolker, B., & Walker, S. (2014). Fitting linear mixed-effects models using lme4. arXiv preprint arXiv:1406.5823.

Bernasconi, F., Grivel, J., Murray, M. M., & Spierer, L. (2010). Plastic brain mechanisms for attaining auditory temporal order judgment proficiency. Neuroimage, 50(3), 1271–1279.

Blakemore, S. J., & Frith, C. (2003). Self-awareness and action. Current opinion in neurobiology, 13(2), 219–224.

Blanke, O., & Metzinger, T. (2009). Full-body illusions and minimal phenomenal selfhood. Trends in cognitive sciences, 13(1), 7–13.

Blanke O. (2012). Multisensory brain mechanisms of bodily self-consciousness. Nature Reviews Neuroscience, 13(8), 556–571.

Blanke, O., Pozeg, P., Hara, M., Heydrich, L., Serino, A., Yamamoto, A., … & Arzy, S. (2014). Neurological and robot-controlled induction of an apparition. Current Biology, 24(22), 2681–2686.

Blanke, O., Slater, M., & Serino, A. (2015). Behavioral, neural, and computational principles of bodily self-consciousness. Neuron, 88(1), 145–166.

Brainard, D. H., & Vision, S. (1997). The psychophysics toolbox. Spatial vision, 10, 433–436.

Canzoneri, E., Di Pellegrino, G., Herbelin, B., Blanke, O., & Serino, A. (2016). Conceptual processing is referenced to the experienced location of the self, not to the location of the physical body. Cognition, 154, 182–192.

Cleeremans A. (2011). The radical plasticity thesis: how the brain learns to be conscious. Frontiers in psychology, 2, 86.

Conover, W. J., & Iman, R. L. (1981). Rank transformations as a bridge between parametric and nonparametric statistics. The American Statistician, 35(3), 124–129.

Ehrsson H. H. (2012). 43 The concept of body ownership and its relation to multisensory integration. The New Handbook of Multisensory Process.

Faivre, N., Salomon, R., & Blanke, O. (2015). Visual consciousness and bodily self-consciousness. Current opinion in neurology, 28(1), 23–28.

Faivre, N., Doenz, J., Scandola, M., Dhanis, H., Ruiz, J. B., Bernasconi, F., … & Blanke, O. (2017). Self-Grounded Vision: Hand Ownership Modulates Visual Location through Cortical β and γ Oscillations. Journal of Neuroscience, 37(1), 11–22.

Faivre, N., Filevich, E., Solovey, G., Kühn, S., & Blanke, O. (2018). Behavioral, modeling, and electrophysiological evidence for supramodality in human metacognition. Journal of Neuroscience, 38(2), 263–277.

Fleming, S. M., Maniscalco, B., Ko, Y., Amendi, N., Ro, T., & Lau, H. (2015). Action-specific disruption of perceptual confidence. Psychological science, 26(1), 89–98.

Fox J. (2003). Effect displays in R for generalised linear models. Journal of statistical software, 8(15), 1–27.

Gajdos, T., Fleming, S., Garcia, M. S., Weindel, G. & Davranche, K. Revealing subthreshold Motor contributions to perceptual confidence. Preprint at https://www.biorxiv.org/content/early/2018/05/25/330605 (2018)

Gallagher S. (2000). Philosophical conceptions of the self: implications for cognitive science. Trends in cognitive sciences, 4(1), 14–21.

Haggard, P., Clark, S., & Kalogeras, J. (2002). Voluntary action and conscious awareness. Nature neuroscience, 5(4), 382–385.

Haggard P. (2017). Sense of agency in the human brain. Nature Reviews Neuroscience, 18(4), 196.

Hara, M., Rognini, G., Evans, N., Blanke, O., Yamamoto, A., Bleuler, H., & Higuchi, T. (2011, September). A novel approach to the manipulation of body-parts ownership using a bilateral master-slave system. In Intelligent Robots and Systems (IROS), 2011 IEEE/RSJ International Conference on (pp. 4664–4669). IEEE.

Kleiner, M., Brainard, D., Pelli, D., Ingling, A., Murray, R., & Broussard, C. (2007). What’s new in Psychtoolbox-3. Perception, 36(14), 1.

Koriat A. (2006). Metacognition and consciousness. Institute of Information Processing and Decision Making, University of Haifa.

Kornbrot D. E. (2006). Signal detection theory, the approach of choice: Model-based and distribution-free measures and evaluation. Perception & Psychophysics, 68(3), 393–414.

Kuznetsova, A., Brockhoff, P. B., & Christensen, R. H. B. (2015). Package ‘lmerTest’. R package version, 2.

Levitt H. C. C. H. (1971). Transformed up-down methods in psychoacoustics. The Journal of the Acoustical society of America, 49(2B), 467–477.

Maniscalco, B., & Lau, H. (2012). A signal detection theoretic approach for estimating metacognitive sensitivity from confidence ratings. Consciousness and cognition, 21(1), 422–430.

Maniscalco, B., & Lau, H. (2014). Signal detection theory analysis of type 1 and type 2 data: meta-d’, response-specific meta-d’, and the unequal variance SDT model. In The cognitive neuroscience of metacognition (pp. 25–66). Springer, Berlin, Heidelberg.

Moore, J. W., & Obhi, S. S. (2012). Intentional binding and the sense of agency: a review. Consciousness and cognition, 21(1), 546–561.

Morey, R. D., Rouder, J. N., Jamil, T., & Morey, M. R. D. (2015). Package ‘BayesFactor’. URL ⟨http://cran.r-project.org/web/packages/BayesFactor/BayesFactor.pdf⟩ (accessed 10.06. 15).

Noel, J. P., Blanke, O., Serino, A., & Salomon, R. (2017). Interplay between narrative and bodily self in access to consciousness: No difference between self-and non-self attributes. Frontiers in psychology, 8, 72.

Orne M. T. (1962). On the social psychology of the psychological experiment: With particular reference to demand characteristics and their implications. American psychologist, 17(11), 776.

Pereira, M., Faivre, N., Iturrate, I., et al. Disentangling the origins of confidence in speeded perceptual judgments through multimodal imaging. BioRxiv, 2018, https://doi.org/10.1101/496877.

Pelli D. G. (1997). The VideoToolbox software for visual psychophysics: Transforming numbers into movies. Spatial vision, 10(4), 437–442.

Pleskac, T. J., & Busemeyer, J. R. (2010). Two-stage dynamic signal detection: a theory of choice, decision time, and confidence. Psychological review, 117(3), 864.

Pouget, A., Drugowitsch, J., & Kepecs, A. (2016). Confidence and certainty: distinct probabilistic quantities for different goals. Nature neuroscience, 19(3), 366–374.

Rausch, M., Müller, H. J., & Zehetleitner, M. (2015). Metacognitive sensitivity of subjective reports of decisional confidence and visual experience. Consciousness and cognition, 35, 192–205.

Rochat P. (2003). Five levels of self-awareness as they unfold early in life. Consciousness and cognition, 12(4), 717–731.

Siedlecka M., Paulewicz B. & Wierzchoń M. But I was so sure! Metacognitive judgments are less accurate given prospectively than retrospectively. Front. Psychol. 7:1–8 (2016).

Singmann, H., Bolker, B., Westfall, J., Højsgaard, S., Fox, J., & Lawrence, M. (2015). afex: Analysis of factorial experiment. R package version 0.13-145.

Stetson, C., Cui, X., Montague, P. R., & Eagleman, D. M. (2006). Motor-sensory recalibration leads to an illusory reversal of action and sensation. Neuron, 51(5), 651–659.

Timmermans, B., Schilbach, L., Pasquali, A., & Cleeremans, A. (2012). Higher order thoughts in action: consciousness as a unconscious re-description process. Philosophical Transcations of the Royal Society B, 367, 1412–1423.

Wenke, D., & Haggard, P. (2009). How voluntary actions modulate time perception. Experimental brain research, 196(3), 311–318.

Wickham H. (2009). ggplot2: elegant graphics for data analysis. Springer New York, 1(2), 3.

Yeung, N., & Summerfield, C. (2012). Metacognition in human decision-making: confidence and error monitoring. Phil. Trans. R. Soc. B, 367(1594), 1310–1321.

